# Neural activity is spatially clustered in motor and dorsal premotor cortex

**DOI:** 10.1101/2022.09.20.508805

**Authors:** Nick G. Chehade, Omar A. Gharbawie

## Abstract

Motor (M1) and dorsal premotor (PMd) cortex are central to arm and hand control in primates. Motor outputs in both areas confer somatotopically organized arm and hand zones. Here, we investigate the spatial mapping between those motor zones and movement-related neural activity to gain insight about functional organization in M1 and PMd. Two macaques reached and grasped while cortical activity was measured with intrinsic signal optical imaging. Activity maps were quantified in relation to microstimulation motor maps from the same hemispheres. Each activity map was comprised of many patches and overlapped surprisingly small portions of the motor map. Functional differences between the patches were inferred from their activity time courses and location within the motor map. We propose that M1 and PMd contain subzones that are preferentially tuned for specific actions. Thus, the spatial dimension of neural activity in frontal motor areas is an important organizing principle of the neural code for movement control.

## Introduction

Motor (M1) and dorsal premotor (PMd) cortex in monkeys are widely used as models for studying cortical control of movement. Both cortical areas contain complete motor representations of the forelimb (Boudrias et al., 2010; Gould et al., 1986; Kwan et al., 1978; Raos et al., 2003). Somatotopy of the forelimb representations varies across studies, but there is consensus that muscles and cortical columns do not have a one-to-one mapping. Instead, an arm or a hand muscle receives corticospinal inputs from a population of cortical columns (Andersen et al., 1975; Rathelot and Strick, 2006). Similarly, a cortical column can influence activity in several muscles (Donoghue et al., 1992; Fetz and Cheney, 1980; Lemon et al., 1987). The convergence and divergence of corticospinal projections means that the same motor output is present in many sites within a forelimb representation. For example, intracortical microstimulation (ICMS) evokes shoulder flexion from many M1 sites where the affiliate corticospinal projections target the same group of arm muscles (Park et al., 2001). But how does a seemingly simple motor map control a rich repertoire of arm and hand actions?

To address this question, we must first understand the spatial relationship between motor map organization and neural activity that support functions. We consider two competing perspectives. (1) Functions are widely encoded throughout motor zones. In this scheme, neural activity that supports arm functions (e.g., reach) would be present throughout motor arm zones. This scheme has support in electrophysiological studies that examined relationships between neural activity, recording site location, and forelimb actions (Rouse and Schieber, 2016; Saleh et al., 2012). Nevertheless, most electrophysiological investigations pool neural activity across recording sites, offering only limited insight into the spatial organization of functions. (2) Functions are clustered within motor zones. In this scheme, neural activity that supports an arm function would be concentrated in subzones of the motor arm zones. Neural activity in adjacent subzones would be tuned for other arm functions. This functional clustering scheme is consistent with maps obtained using long train ICMS (500 ms), which approximates the duration of natural actions (Graziano et al., 2002). The same motor mapping parameters revealed functional subzones in both frontal and parietal cortical areas (Cooke and Graziano, 2004; Gharbawie et al., 2011; Graziano et al., 2002; Stepniewska et al., 2005).

Adjudicating between the two schemes can be facilitated with neuroimaging, which provides uninterrupted spatial sampling and retains the spatial dimension of the recorded activity. The necessary implementation here is to image entire forelimb representations during arm and hand actions then quantify the spatial organization of movement-related cortical activity. Intrinsic signal optical imaging (ISOI), 2-photon imaging, and fMRI have been successfully used in measuring cortical activity related to arm and hand actions in monkeys (Ebina et al., 2018; Friedman et al., 2020a; Kondo et al., 2018; Nelissen and Vanduffel, 2011). The mesoscopic field-of-view in ISOI is particularly well-suited for the size of the forelimb representations in M1 and PMd. Moreover, ISOI affords high spatial resolution and contrast without extrinsic agents (e.g., GCaMP, dyes, MION). These reasons motivated us to use ISOI to measure M1 and PMd activity in head-fixed macaques engaged in an instructed reach-to-grasp task (Fig. 1A-D). In the same field-of-view, we used ICMS to map the forelimb representations and surrounding territories. We quantified the overlap between ISOI maps and forelimb motor maps for insight into the spatial organization of task-related cortical activity. In a distributed functional organization, we would expect ISOI to report task-related activity throughout the forelimb representations in M1 and PMd (Fig. 1B top). In contrast, in a clustered functional organization, ISOI would report task-related activity in subzones of the forelimb representations (Fig. 1C bottom).

**Figure 1.**
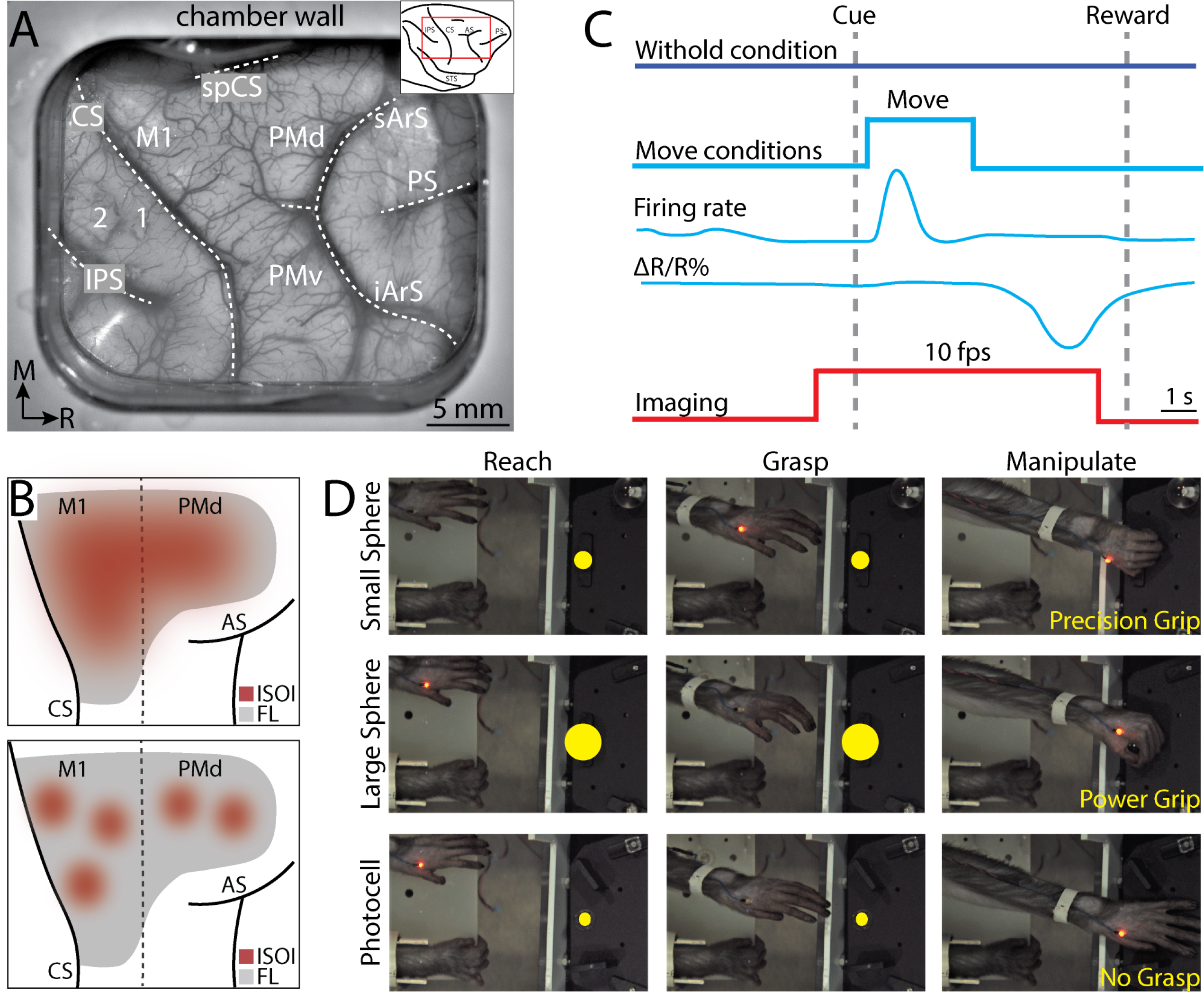
Intrinsic signal optical imaging during forelimb arm and hand actions. (A) Chronic recording chamber provides access to motor and somatosensory areas in the right hemisphere. Native dura was replaced with transparent membrane. Dashed lines mark: central sulcus (CS), intraparietal sulcus (IPS), and arcuate sulcus (AS). Inset shows approximate location of chamber (red rectangle). (B) Schematic of potential results. Dotted line separates M1 from premotor areas. Forelimb representation is gray. Red patches are clusters of pixels that darkened (i.e., negative reflectance) after task-related increase in neural activity. (Top) *H_0_*: pixels darken throughout the forelimb representation. (Bottom) *H_1_*: pixels darken in spatial clusters within the forelimb representation. (C) Relative timing in task conditions. Blue pulse indicates movement period, whereas there was no movement in the withhold condition. Increase in neural firing coincides with movement and precedes reflectance change (ΔR/R%) measured with ISOI. Red pulse depicts ISOI acquisition in both conditions (10 frames per second). (D) Still frames from 3 phases (columns) in the 3 movement conditions (rows). Task was performed with the left forelimb and the right forelimb was restrained.

## Methods

### Animals

The right hemisphere was studied in two male macaque monkeys (*Macaca mulatta*). Monkeys were 6-7 years old and weighed 9-11 kg. All procedures were approved by the University of Pittsburgh Animal Care and Use Committees and followed the guidelines of the National Institutes of Health guide for the care and use of laboratory animals.

### Head post and recording chamber

After an animal acclimated to the primate chair and training environment, a head-fixation device was secured to the occipital bone and caudal parts of the parietal bone. Task training with head-fixation started after ∼1 month (monkey G) or ∼9 months (monkey S) and lasted for ∼22 months. A craniotomy was then performed for a chronic recording chamber (30.5 x 25.5 mm internal dimensions) over motor and somatosensory cortical areas. The dura was resected and replaced with a transparent silicone membrane for visualization of cortical blood vessels and landmarks (Fig. 1A).

### Reach-to-grasp task

Monkeys performed a reach-to-grasp task while head fixed in a primate chair. The left forelimb was used, and the right forelimb was secured to the waist plate. The task apparatus was positioned in front of the animal. A stepper motor rotated the carousel in between trials to present a target ∼200 mm from the start position of the left hand. Targets were presented in the same location to encourage consistent reach trajectories. Task instruction was provided with LEDs mounted above the target. Photocells were embedded in multiple locations within the apparatus to monitor hand location and target manipulation. An Arduino board (Arduino Mega 2560, www.arduino.cc) running a custom script (1 kHz) controlled task parameters, timing, and logged the monkey’s performance on each trial. The task involved three kinds of conditions.

#### 1. Reach-to-grasp

In a successful trial, the animal had to reach, grasp, lift, and hold a sphere. Small (12.7 mm diameter) and large (31.8 mm diameter) spheres were used to motivate precision and power grips, respectively (Fig. 1D). Spheres were attached to rods that moved in a vertical axis only. To initiate a trial, an animal placed its left hand over a photocell embedded in the waist plate. Covering the photocell for 300 ms turned on an LED, which indicated the start of the trial. Holding this start position for 5000 ms triggered the Go Cue, which was a blinking LED. The animal had 2400 ms (monkey G) or 2550 ms (monkey S) to reach, grasp, and lift the sphere; time limits were also set for each phase. Lifting the sphere by 1500 mm turned the blinking LED solid, which signaled the end of the lift phase. Maintaining the lifted position for 1000 ms turned off the LED, which instructed the animal to release the object and return its hand to the start position within 900 ms. Maintaining the start position for 5000 ms triggered a tone and LED blinking. After an additional 2000 ms in the start position the trial was considered successful; tone and LEDs turned off and water reward was delivered. The animal could not initiate a new trial for another 3000 ms. Failure to complete any step within the allotted time window resulted in an incorrect trial signaled by a 1500 ms tone and a 5000 ms timeout in which the apparatus was unresponsive to the monkey’s actions. After the timeout, a new trial could be initiated with hand placement in the start position.

#### 2. Reach-only condition

The target was a photocell embedded into the surface of the carousel. The photocell was visible to the monkey but was not graspable. The Go Cue prompted the monkey to reach and place its hand over the photocell. The hand had to cover the photocell for 220 ms (monkey G) or 320 ms (monkey S). All other task rules and steps were identical to the reach-to-grasp condition.

#### 3. Withhold condition

In a successful trial, the monkey had to maintain its hand in the start position for ∼10 s. Trial initiation was identical to other conditions. Holding the start position for 5000 ms triggered the Withhold Cue, which was distinctly different from the Go Cue in the movement conditions. Maintaining the start position for another 2800 ms triggered a tone and LED blinking. After an additional 2000 ms in the start position the trial was considered successful and rewarded. Removing the hand from the start position at any time resulted in an incorrect trial and the same consequences described in a failed reach-to-grasp trial.

Conditions were presented in an event-related design (1 successful trial/condition/block). Condition order was randomized across blocks. We structured the trials and the inter-trial interval so that the hand would remain in the start position for ∼13 s in between trials. Thus, in a successful trial from a movement condition, the arm and hand remained still in the start position for ∼12.3 s and then moved for 0.8-1.8 s. This relatively long period without movement was useful for relating intrinsic signal changes to movement onset, which is a considerably more rapid change in behavior. In the withhold condition, the arm and hand were in the start position for ∼10 s and no movement was allowed.

### Muscle activity

EMG activity was recorded from 7 forelimb muscles on select sessions. Before the task started, two stainless steel wires (27 gauge, AM Systems) were inserted percutaneously into each muscle. Three arm muscles were targeted (1) deltoideus, (2) triceps brachii, and (3) biceps brachii. Four extrinsic hand muscles were targeted (1) extensor carpi radialis brevis, (2) flexor carpi radialis, (3) extensor digitorum 4-5, and (4) flexor digitorum superficialis. EMG signals were bandpass filtered at 15-350 Hz and digitized at 2000 Hz (Scout Processor, Ripple Neuro). Recorded signals were segmented into trials and their power spectrum density was estimated with a discrete Fourier transform (MATLAB *fft* function, Natick, MA). Trials with power >7 μV^2^ in the 1-14 Hz range were presumed to have artifact and were excluded from further analysis. EMG signals were rectified and smoothed with a 100 ms sliding window (MATLAB *filtfilt* function). Finally, for each muscle in each movement condition, the average signal was computed across trials and sessions (308-595 trials/muscle).

### Joint kinematics

Forelimb joints were tracked in 3D on select sessions. Before the task started, 6 LEDs were secured to sites on the left forelimb to track up to two joints/session. A motion capture system (Impulse X2, Phasespace) outfitted with six cameras recorded LED positions (480 Hz) and logged x, y, z coordinates. For each tracked joint, LEDs were configured to form imaginary vectors or planes. For example, to track the elbow, 3 LEDs were placed in a triangular formation on the upper arm (i.e., parallel to humerus) and another 3-LED formation was placed on the forearm (i.e., parallel to ulna). Elbow flexion/extension was calculated as the angle between the vector aligned with the upper arm and the vector aligned with the forearm. Pronation/supination was calculated as the angle between the normal of the plane of the upper arm and the normal of the plane of the forearm. A similar approach was adopted for (1) shoulder flexion/extension, (2) shoulder abduction/adduction, (3) wrist flexion/extension, and (4) digits flexion/extension. Time series of LED coordinates were segmented into trials. Trials were excluded from analyses if LED positions were not logged for >2% of trial duration. Such dropouts were typically due to LED occlusion by the forelimb, primate chair, or grasp apparatus. Kinematic profiles were calculated trial-by-trial and then averaged for each condition (143-405 trials/degree of freedom).

### Motor map

We used intracortical microstimulation (ICMS) to map the somatotopic organization of frontal motor areas. In monkey S, all sites (n=158) were investigated with a microelectrode in dedicated motor mapping sessions. We used the same approach in >50% of sites (n=118) in monkey G. The remaining sites (n=99) were mapped with a linear electrode array at the end of electrophysiological recordings that will be presented in a separate report. In the dedicated motor mapping sessions, the monkey was head-fixed in the primate chair and sedated (ketamine, 2.0– 3.0 mg/kg, i.m., every 60-90 minutes). This mild sedation reduced voluntary movements but did not suppress reflexes or muscle tone. A hydraulic microdrive (Narishige MO-10) connected to a customized 3-axis micromanipulator was attached to the recording chamber for positioning a tungsten microelectrode [250 μm shaft diameter, impedance = 850 ± 97 kΩ (mean ± SD)] or a platinum/iridium microelectrode [250 μm shaft diameter, impedance = 660 ± 153 kΩ (mean ± SD)]. A surgical microscope aided with microelectrode placement in relation to cortical microvessels. The microelectrode was in recording mode at the start of every penetration.

Voltage differential was amplified (10,000×) and filtered (bandpass 300–5000 Hz) using an AC Amplifier (Model 2800, AM Systems). The signal was passed through a 50/60 Hz noise eliminator (HumBug, Quest Scientific Instruments Inc.) and monitored with an oscilloscope and a loudspeaker. As the electrode was lowered, the first evidence of neural activity was considered 500 μm below the pial surface. The microelectrode was then switched to stimulation mode and the effects of ICMS were tested at four depths (500, 1000, 1500, 2000 μm). Microstimulation trains (18 monophasic, cathodal pulses, 0.2 ms pulse width, 300 Hz) were delivered from an 8-Channel Stimulator (model 3800, AM Systems). Current amplitude, controlled with a stimulus isolation unit (model BSI-2A, BAK Electronics), was increased until a movement was evoked (max 300 μA). Threshold for a stimulation depth was the current amplitude that evoked movement on 50% of stimulation trains.

One experimenter controlled the location and depth of the microelectrode. A second experimenter, blind to microelectrode location, controlled the microstimulation. Both experimenters inspected the evoked movement and classified it according to joint (i.e., digits, elbow, etc.) and movement type (flexion, extension, etc.). The overall classification for a penetration included all movements evoked within 30% of the lowest threshold across depths. The location of each penetration (500-1000 μm apart) was recorded in relation to cortical microvessels. Color-coded maps were generated from this data using a voronoi diagram (MATLAB *voronoi* function) with a maximum tile radius of 750 μm (Fig. 2B & 2F). The rostral border of M1 was marked to separate sites with thresholds <30 μA from higher threshold sites (Fig. 2C & 2G).

**Figure 2.**
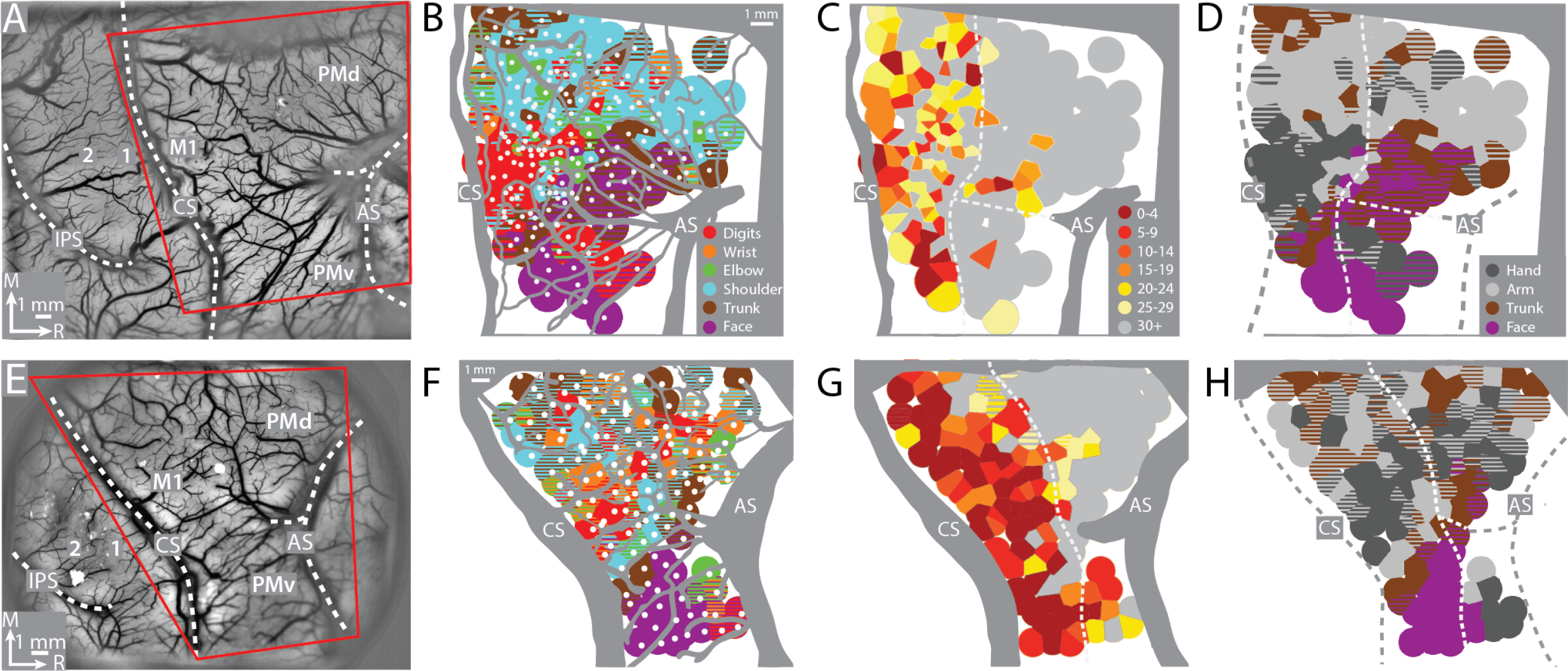
Motor map organization in M1 and PMd. (A-D) Right hemisphere of monkey G. (A) Cropped image of chronic chamber. Blood vessels and cortical landmarks are visible through the transparent membrane. Red outline is the field-of-view in subsequent panels and figures. (B) Major blood vessels and chamber walls are masked in gray. White dots are intracortical microstimulation (ICMS) sites (n = 218). Voronoi tiles (0.75 mm radius) are color-coded according to ICMS-evoked movement. Striped tiles represent dual movements. (C) Same motor map from (B) colored according to current amplitude (μA) for evoking movements. Border between M1 and premotor cortex is drawn at the transition from low (<30 μA) to high (≥30 μA) current thresholds. (D) Same motor map as in (A), but wrist and digit sites are classified as hand, and shoulder and elbow sites are classified as arm. (E-H) Same as top row, but for right hemisphere of monkey S. Motor map has 158 ICMS sites.

The mapping approach was similar for sites stimulated with a linear electrode array (32 or 24 channels, 15 μm contact diameter, 100 μm inter-contact distance, 210-260 μm probe diameter, V-Probe, Plexon). Each site was mapped 2.0-2.5 hours after the array was inserted into cortex, which was also the end of electrophysiological recordings from the penetration. Only 1 site was mapped per session. Microstimulation trains were identical to the ones used with the microelectrode but were delivered from a head-stage that supports both recording and stimulation (Nano2+Stim, Ripple Neuro). Current amplitude was controlled from the Trellis Software Suite (Ripple Neuro). Channels were stimulated one at a time and every other channel was used. One experimenter controlled current amplitude and classified the evoked movements.

### Intrinsic signal optical imaging

The field-of-view (FOV) was illuminated with 2-3 independently controlled red LEDs (630 nm wavelength). Camera frames 768×768 pixels (monkey G) or 1080×1310 pixels (monkey S) were captured with a 12-bit CMOS sensor (Photon Focus, Lachen, Switzerland). The tandem lens combination achieved a FOV of 15×15 mm or 22×26 mm, both at 20 μm^2^/pixel. Frames were temporally averaged from 100 frames/s to 10 frames/s then saved. Image acquisition and parameters were controlled with an Imager 3001 system (Optical Imaging Ltd, Rehovot, Israel). In every trial, imaging started 1 s before Cue onset and lasted for 7-10 s. Every imaging session included 35-60 correct trials/condition. Analyses were based on correct trials only. For spatial reference, a high contrast image of the cortical surface was captured with green illumination (528 nm wavelength) at the start of every session.

### Image processing

Image processing was conducted on individual imaging sessions. Data frames were rigid aligned in x, y coordinates (MATLAB *estimateGeometricTransform* function) to a reference frame from the middle of the session. A trial was excluded if any frame was out of register by >10 pixels (i.e., >200 μm). In the remaining trials, the first frame was subtracted from all frames to increase signal-to-noise ratio. The first frame was the first frame in a trial (−1.0 second from Cue) or an average of the initial four frames in a trial (−1.0 to −0.7 s from Cue). First frame subtraction converted pixel values to reflectance change in relation to baseline. Trials from the same condition were averaged to generate a mean time series. Average frames were processed with a high-pass median filter (kernel = 250 or 550 pixels) to correct uneven illumination and residual motion artifact. A low-pass Gaussian filter (kernel = 5 or 15 pixels) was used for smoothing. To enhance contrast, pixel values were clipped in relation to the median pixel value.

### Reflectance change time course

Time course profiles were examined in average time series. Image processing was same as above, but clipping was excluded. Time courses were generated for regions-of-interest (ROI) that were placed in M1 and PMd. Pixel values within an ROI were averaged to obtain 1 value per frame. Pixels that overlapped blood vessels were excluded.

### Activity maps

Pixel darkening is a lagging indicator of increased neural activity. To identify pixels that darkened in response to instructed movement, we compared frames from the end of movement (i.e., movement frames) to frames acquired before movement onset (i.e., baseline frames). Comparisons were conducted on individual sessions. Every trial contributed 1 movement frame and 1 baseline frame. A baseline frame was an average of 5 or 9 data frames acquired −1.0 s from Cue. A movement frame was an average of 5 data frames acquired +4.0 s from Cue, which was also 1.0-1.5 s after completion of task-related movements. Reflectance change time courses guided our selection of movement frames. A two-sample t-test was then conducted pixel-by-pixel to compare values between movement frames and baseline frames. Right tail values (p<0.0001) indicated that the affiliate pixels brightened significantly in movement frames as compared to baseline frames. Left tail values (p<0.0001) indicated that pixels darkened significantly in movement frames as compared to baseline frames. We flagged the darkened pixels to generate a *thresholded frame* for the session.

Threholded frames from different sessions were aligned to a common reference, which was a high contrast image of the cortical surface. For every imaging session, we marked 25-60 points that were evident in the microvessel patterns present in that session and in the common reference. These points were used to construct a mesh grid for the reference and session images. Non-rigid transformation (multilevel B-spline approximation) was then applied to fit the session mesh grid to the reference mesh grid (Koon, 2008; Lee et al., 1997; Rueckert et al., 1999). For each session, the transformation matrix was used to co-register the thresholded frame to the common reference. From the stack of thresholded frames, pixels flagged in ≥ 50% of sessions were synthesized into an *activity map*.

### Activity map quantification

Activity maps were quantified in relation to the motor map, which was already registered to the common reference. Pixels in activity maps were counted in each somatotopic zone (e.g., M1 hand, PMd arm, etc.). Pixels that overlapped Voronoi tiles with multiple representations (e.g., M1 arm and M1 hand) were counted once per representation. Pixel counts were converted into surface area (mm^2^) and compared across task conditions.

### k-medoids clustering

We used k-medoids to cluster ROIs with similar time course profiles. ROIs were a grid of cells (1 cell = 15×15 pixels; 0.27×0.27 mm) that covered the FOV. Time courses were generated for all cells in each imaging session. Spatially matched cells were co-registered (non-rigid transform described above) across sessions and the average time course was calculated for each cell. Average time courses were smoothed with a 200 ms sliding window (MATLAB *movmean* function) and then used as input for clustering.

The clustering algorithm (MATLAB *kmedoids* function) operates on a feature space, which was defined here as a matrix of cells (n=1700 or n=2249) x time courses (70 values/time course). The distance metric of the clustering algorithm was the max cross-correlation coefficient (MATLAB *xcorr* function) for each pair of cells. Coefficients were considered from every possible time lag (141 lags, −7.0 to 7.0 s in 100 ms steps) and were normalized by dividing by the total number of lags. Negative coefficients were set to zero. The *kmedoids* function was run with the number of clusters set from 1 to 10. In every run, we first calculated the *within cluster sum,* which is the sum of distances between every cell in the cluster and the medioid of that cluster. Then we calculated the *total distance,* which is the total sum of the within cluster sums. *Total distance* was then fed into the triangle threshold method to determine the optimal number of clusters. To that end, *total distance* (y-axis) was plotted against the number of clusters (x-axis). The tops of the largest and smallest bars were then connected with a line. From this line, orthogonal lines were projected to the top of every bar. The longest of those orthogonal line identified the optimal number of clusters (Panneton, 2010; Zack et al., 1977).

## Results

Two monkeys performed an instructed forelimb task that consisted of 4 conditions: (1) reach-to-grasp with precision grip, (2) reach-to-grasp with power grip, (3) reach-only, and (4) withhold (Fig. 1D). We used intrinsic signal optical imaging (ISOI) to measure cortical activity during task performance (Fig. 1). Activity in M1 and PMd was spatially quantified in relation to motor maps obtained with intracortical microstimulation (ICMS) from the same recording chambers. We used the results to evaluate competing perspectives about the functional organization of forelimb representations in M1 and PMd (Fig. 1B).

### Consistent somatotopy in M1 and PMd

To find the forelimb representations, we used ICMS to map frontal cortex from central sulcus to arcuate sulcus (Fig. 2A & 2E). The general organization of the motor map was consistent between monkeys (Fig. 2B-D & 2F-H). Most ICMS sites evoked forelimb movements (Fig. 2B & 2F). We marked the rostral border of M1 such that (1) it was 3-5 mm from the central sulcus, and (2) separated high threshold sites (>30 μA) from low threshold sites (Fig. 2C & 2G). To simplify the motor map, site classifications were consolidated into broader categories (e.g., elbow and shoulder became arm). The simplified maps (Fig. 2D & 2H) showed that the forelimb representation was flanked medially by trunk zones and laterally by trunk and face zones. Within the M1 forelimb representation, the main hand zone (i.e., digits and wrist) was surrounded by an arm zone, or an arm and trunk zone. This nested organization is consistent with previous maps from macaque monkeys (Kwan et al., 1978; Park et al., 2001; Sessle and Wiesendanger, 1982). The organization of the forelimb representations was less clear in PMd and PMv, which could have been related to higher thresholds, less extensive mapping, as well as more overlap between arm and hand zones than in M1 (Boudrias et al., 2010).

### Movement-related activity is clustered in M1 and PMd

Neural activity drives a hemodynamic response that is detectable as reflectance change in ISOI. Under red light illumination (630 nm wavelength), which was used here, negative reflectance (i.e., pixel darkening) is a lagging indicator of increased neural activity (Fig. 1C). Thus, in movement conditions, pixel darkening reports locations where neural activity increased for movement execution, or movement planning, or both. We refer to clusters of darkened pixels as “domains”. If task-related neural activity increases throughout the forelimb representations, then we could expect a large domain to fill the field-of-view (Fig. 1B top). But if neural activity is more spatially confined, then we could expect multiple, smaller, domains (Fig. 1B bottom).

First, we examined reflectance change in M1 and PMd in average time series from a representative session (36 trials/condition). In the power grip condition (Fig. 3A), there was no reflectance change from baseline to movement onset (−1.0 to +0.5 s from Cue). During reach, grasp, and hold (+1.0 to 3.0 s from Cue), pixels in the center of the FOV brightened (cool colors) and pixels near the perimeter darkened (warm colors). Once the hand returned to the start position (i.e., movement completion), pixels in the center of the FOV started to darken and domains in M1 and PMd began to form (+3.5 to +3.8 s from Cue). Domains continued to darken and expand in the remaining frames. Nevertheless, even at peak expansion and intensity (+5.0 to +6.0 s from Cue), domains remained spatially separable within M1 and between M1 and PMd. For contrast, Fig. 3B shows an average time series from the withhold condition where the hand remained in the start position. Here, pixel darkening was limited to the perimeter as most of the FOV brightened (Fig. 3B, cyan pixels). Thus, domains in Fig. 3A were likely locations with movement-related increase in neural activity.

**Figure 3.**
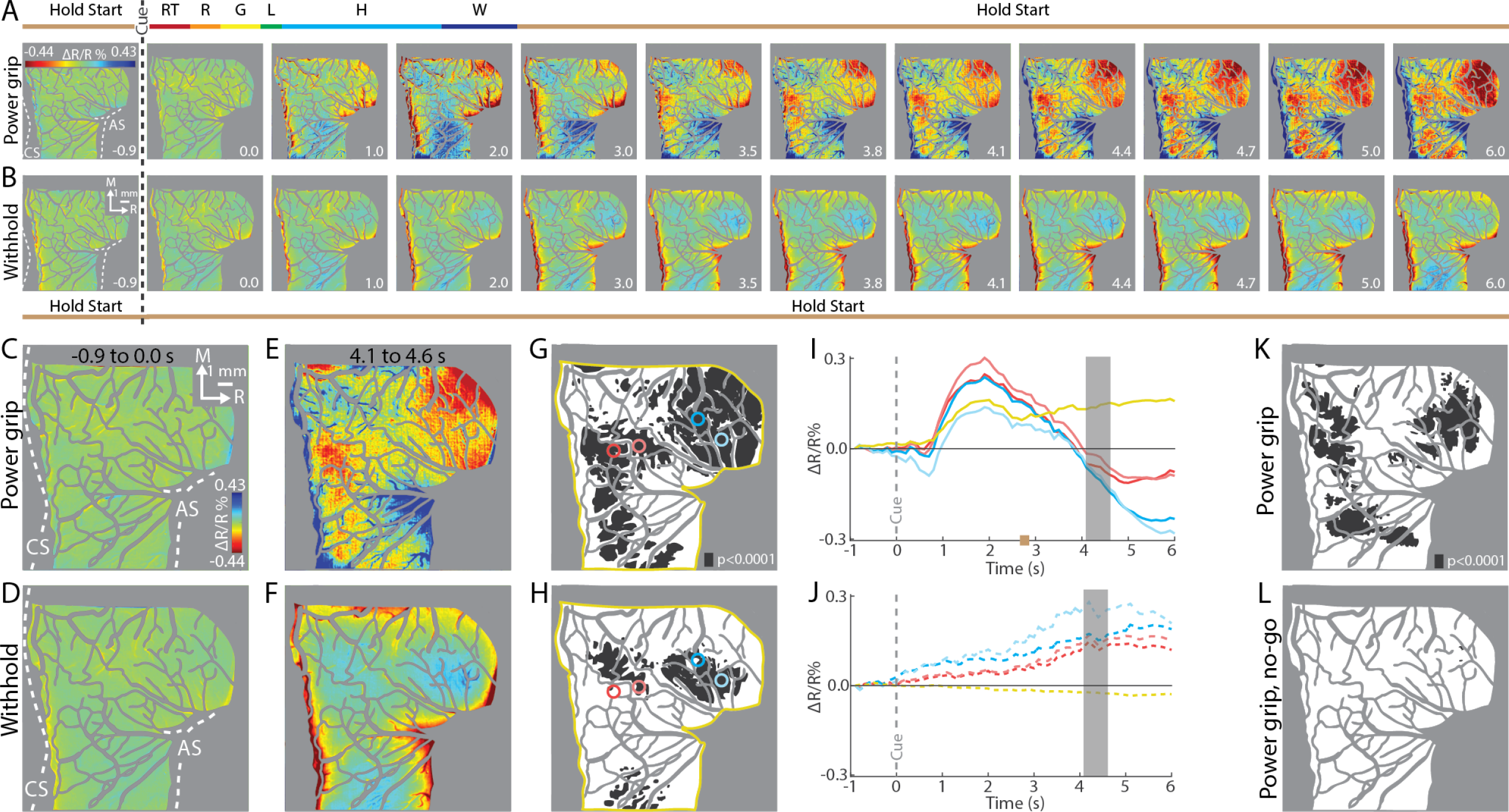
ISOI detects movement-related activity in M1 and PMd. (A-B) Mean time series from a representative session (36 trials/condition, monkey G). (A) Timeline of average trial from the power grip condition. Hold start = hand in start position. Other phases include reaction time (RT), reach (R), grasp (G), lift (L), hold (H), and withdraw (W). Select frames with time (seconds from cue onset) in bottom right. Reflectance change was clipped to median ± 0.3 SD pixel value across frames. Clusters of pixels started to darken (hot colors) at 3.5 s and peaked in size and intensity near the last frame. (B) Matching frames from the withhold condition. Color scale is same as (A). (C-D) Mean baseline frames from the power grip and withhold conditions. (E) Mean of 5 frames captured after movement completion (+4.1 to +4.6 s from Cue) in the power grip condition. (F) Temporally matched mean frame from the withhold condition. (G) Gray pixels were darker (t-test, left tail, p < 0.0001) in (E) than in (C). Colored circles (0.36 mm radius) and yellow outline are regions-of-interest (ROIs). (H) Grey pixels were brighter (t-test, right tail, p < 0.0001) in (F) than in (D). (I) Time courses of reflectance change in the power grip condition. Line colors match the ROIs. Negative values indicate pixel darkening. Grey rectangle matches time window in (E). Brown mark on x-axis is time of movement offset (mean ± SD). (J) Same as (I), but for withhold condition. (K) Activity map from reach-to-grasp condition (2 sessions, 159 trials). Grey pixels darkened after movement as compared to baseline (t-test, left tail, p < 0.0001). (L) Activity map from no-go condition (2 sessions, 159 trials). Only a few pixels darkened in PMd.

We averaged select frames from each time series to examine cortical activity from two phases. For the baseline phase, we averaged frames −1.0 to 0 s from Cue. Average baseline frames showed no reflectance change and were indistinguishable between conditions (Fig. 3C-D). In contrast, differences were apparent between average frames from the post-movement phase (+4.1 to +4.6 s from Cue, Fig. 3E) and the matching period in the withhold condition (Fig. 3F). To threshold Fig. 3E, we flagged pixels that darkened in that frame as compared to the baseline frame (t-test, left tail, p<0.0001, Fig. 3G). For contrast, we thresholded Fig. 3F but focused on pixels that brightened to capture the predominant reflectance change. We therefore flagged pixels that brightened in the withhold frame as compared to the baseline frame (t-test, right tail, p<0.0001, Fig. 3H). Thus, thresholded frames summarized reflectance change in the movement and withhold conditions and indicated stark differences between them (Fig. 3G-H).

Next, we examined the time course of reflectance change in both conditions. We placed four regions-of-interest (ROI) over pixels that darkened in the movement condition (Fig. 3G, colored circles). The two caudal ROIs were in M1, and the two rostral ROIs were in PMd. Time courses were generally consistent across ROIs (Fig. 3I). Positive reflectance started to increase 200-300 ms after movement onset (+0.8 s from Cue) and peaked during the hold phase (+2.0 s from Cue) when the object was grasped and maintained in the lifted position. Then, negative reflectance increased and peaked ∼2 s after task-related movement ended (+5.0 to +6.0 s after Go Cue). The positive–negative sequence will be discussed later in relation to triphasic time courses (negative-positive-negative) established for sensory cortex (Chen-Bee et al., 2007; Sirotin and Das, 2009). For reference, an ROI over the entire FOV (Fig. 3G, yellow outline) had a time course with a positive peak during movement and no change afterwards (Fig. 3I). Time courses were generated from the same ROIs in the withhold condition (Fig. 3H). For the ROIs in M1 and PMd, reflectance drifted positive and lacked distinct peaks. Reflectance in the entire FOV remained near zero. Time courses in Figs. 3I-J indicate that pixel darkening occurred only in the movement condition and only in specific cortical locations.

### Pixel darkening in M1 and PMd is locked to movement

Our next objective was to examine whether movement execution was necessary for the pixel darkening observed in the movement condition. We addressed this point in three experiments. In *Experiment 1*, we added conditions in which the monkey did not move but observed an experimenter perform the task instead. First, the monkey completed blocks of trials with randomized presentation of reach-to-grasp and reach-only conditions (performed trials).

Next, the monkey observed the experimenter perform blocks of trials of the same conditions (observed trials). The primate chair was closed off in those blocks to prevent the monkey from reaching. Cues, timing, and reward schedule were consistent between performed and observed trials. Time courses were measured from ROIs in M1 and PMd (Fig. S1A). In M1, time courses had a negative peak in the performed conditions (Fig. S1B) but remained near baseline in the observed conditions. PMd time courses also had negative peaks in the performed conditions, but they were earlier than in M1. For the observed condition, the small negative peak in the PMd time course stood out from the matching M1 time course. Results from *Experiment 1* indicated that pixel darkening in both M1 and PMd was driven primarily by movement execution. Pixel darkening related to movement preparation, or action processing, was limited to PMd.

In *Experiment 2*, one monkey performed reach-to-grasp trials interleaved with no-go trials. In a no-go trial, the go Cue and preceding steps were no different from a reach-to-grasp trial. But 250 ms after the go Cue (i.e., during reaction time), a no-go Cue (LED and tone) instructed the monkey to abort. Thus, in a correct no-go trial, the monkey had to (1) start reaching, (2) stop its arm before the midpoint between the start position and the target, and (3) return its hand to the start position. Correct trials were typically achieved with the monkey releasing the start position to reach and then immediately returning its hand to the start position. Movements were therefore truncated but not inhibited. We reasoned that no-go trials would minimize movement-related cortical activity without interfering with internal processes that precede movement (e.g., movement preparation). In two representative sessions, clusters of pixels darkened in movement frames as compared to baseline frames (Fig. 5K, t-test, p<0.0001). In contrast, no significant pixel darkening was observed in the no-go condition. *Experiment 2* therefore showed that significant pixel darkening was contingent on movement execution.

Although reach-to-grasp and no-go trials had identical starts, the monkey might still have correctly anticipated the condition before Cue onset. This motivated us to manipulate the timing of the go Cue in *Experiment 3*. Thus, in select sessions, Cue onset was set to 1, 2, or 3 s from trial initiation. We reasoned that temporal shifts in movement onset would lead to predictable shifts in pixel darkening. ROIs were placed in PMd and M1 to examine reflectance change time courses (insets in Figs. 4A-B). Indeed, negative peaks in both PMd and M1 were locked to Cue onset (Fig. 4A-D). This temporal relationship was also evident in the spatial development of the domains (Fig. 4E). For each Cue condition, we generated 3 thresholded frames from 3 time-windows (Fig. 4A, gray). In a thresholded frame, pixels were flagged if they darkened in the time window as compared to the baseline frame (Fig. 4E, t-test; n=44 trials). The first row shows that domains were largest in the first column for the earliest Cue onset. The same spatial pattern was present in the middle column of the second row and in the last panel of the third row. Thus, *Experiment 3* showed that shifting movement onset led to systematic temporal shifts in peak domain size but had no bearing on the spatial organization of the domains. Collectively, *Experiments 1-3* confirm that the present domains were linked to movement.

**Figure 4.**
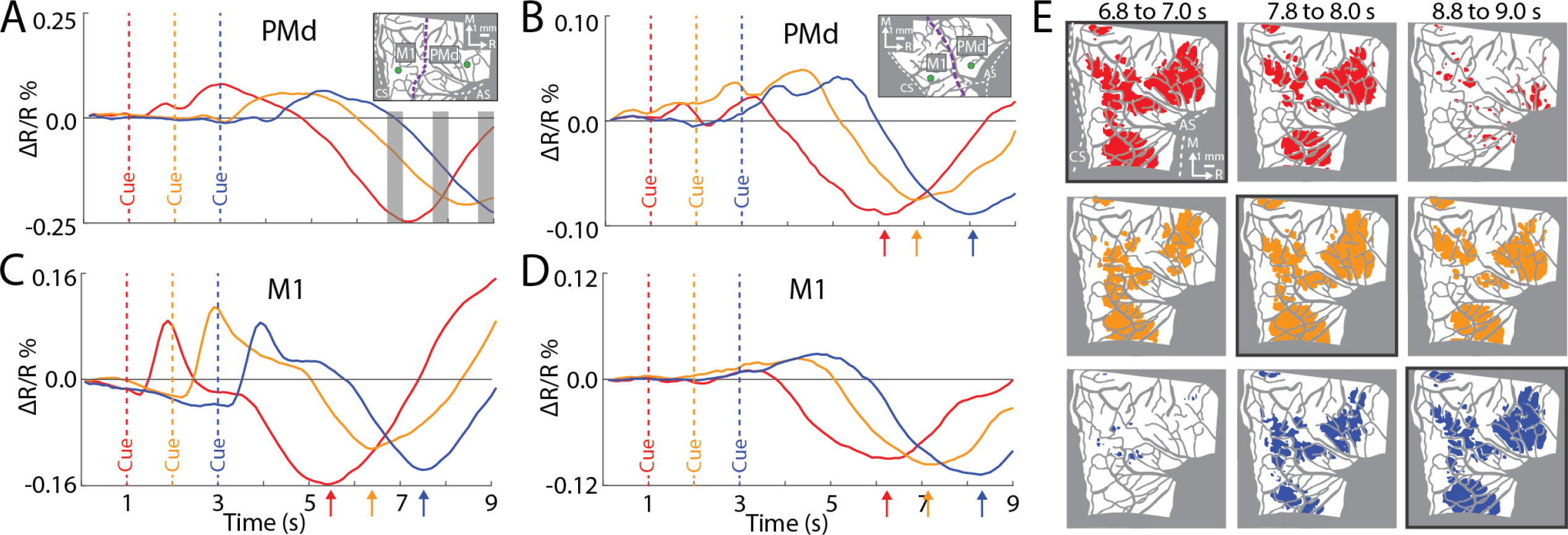
Reflectance change is locked to movement. (A) Time course of reflectance change in PMd (1 session, 44 trials/condition, monkey G). Inset shows green ROIs (0.36 mm radius). Line plot colors match cue condition (dashed line). Grey rectangles correspond to frame ranges in (E). (B) Same as (A), but average of 2 sessions (95 trials/condition, monkey S). Arrows point to negative peaks. (C-D) M1 time courses from the same sessions in (A-B). (E) Colored pixels darkened in the movement frame as compared to baseline (t-test, p<0.0001). Pixel colors match cue condition (1 condition/row). Black borders flag frames based on averaging from the time window: +5.8 to +6.0 s from Cue.

### Consistent features of cortical activity

To identify the most consistent patterns of reflectance change for each condition, we generated average time series from 8 sessions (236-432 trials/condition). Fig. 5A summarizes the average time series for the precision grip condition (monkey G). Animating the time series next to a representative behavioral trial crystalized two spatiotemporal features (Video 1). First, most pixel darkening occurred after movement completion. Second, the darkest pixels were concentrated in spatial clusters that shifted and expanded across the time series (Video 1, grey outline). The same features were present in the other movement conditions (not shown).

**Figure 5.**
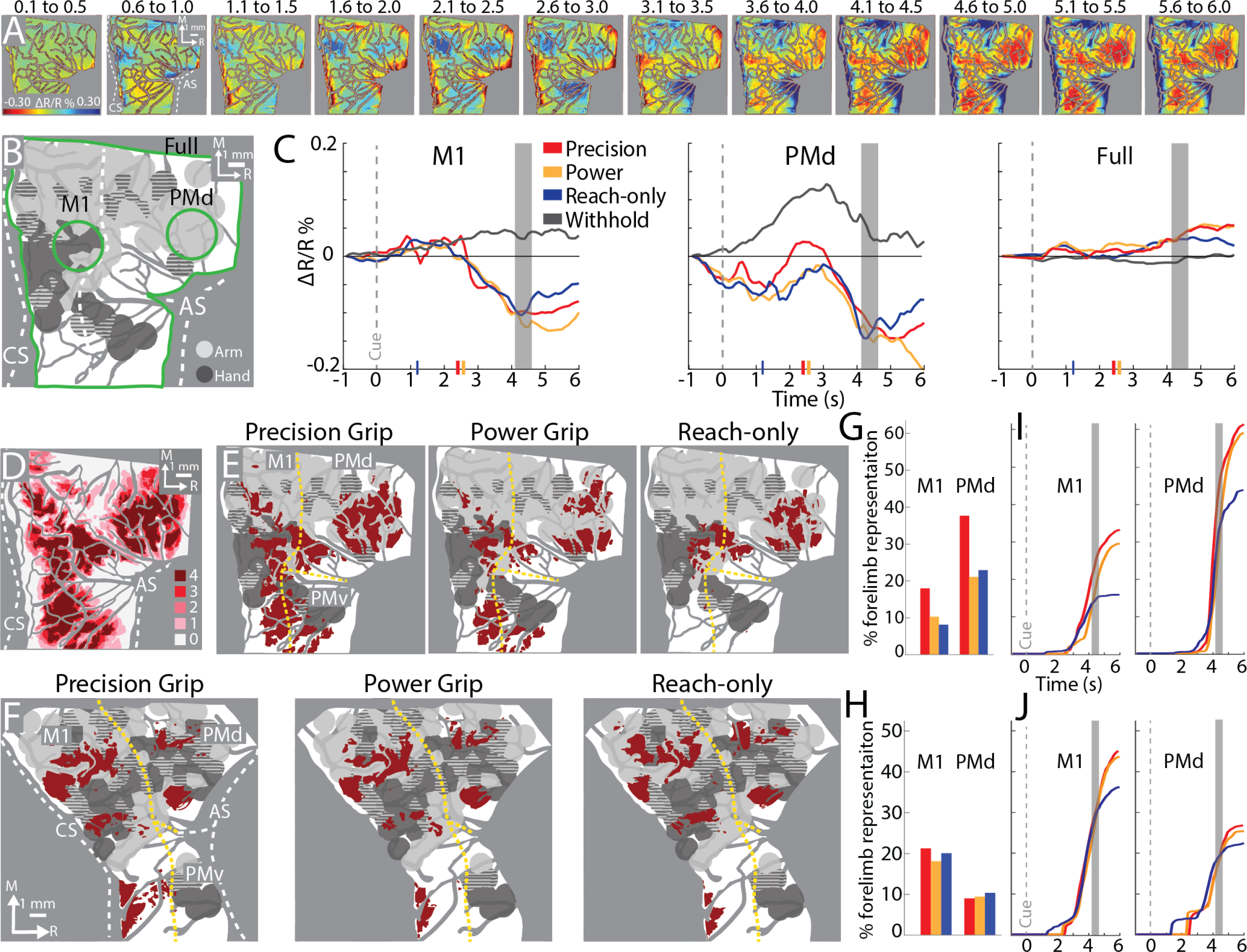
Average time courses and activity maps for different conditions. (A) Average time series for precision grip (8 sessions, 404 trials, monkey G). Each frame is the median of five successive frames and time is seconds from cue onset. Reflectance change was clipped to median ± SD pixel value across frames. Domains started to form at 3.1-3.5 s. (B) Green ROIs in M1 (2.7 mm diameter), PMd, and the full FOV. (C) Median time courses measured from ROIs in (A). Grey rectangles mark the frame range for (D-E). Markers on x-axis match legend and depict movement offset times (mean ± SD). (D) Stack of thresholded frames for the precision grip condition. Pixel intensity reflects number of sessions with darkening in the movement frame as compared to the baseline frame (t-test, p<0.0001). (E) Activity map thresholded to show pixels that darkened in ≥50% of sessions (i.e., ≥4 out of 8 sessions). Dashed yellow lines mark cortical borders. (F) Same as (E), but for monkey S. (G-H) Percentage of forelimb representations occupied by activity maps in (E-F). Bar colors match conditions in (C). (I-J) Same variables in (G-H) expressed as cumulative sums over time. Activity maps were generated for every time point (0.1 s); t-test (p<0.0001) comparison between 5-frame average (centered on time point) and the baseline frame. Frames acquired during movement were excluded to ensure that motion artifact did not contribute to the results.

We also examined time courses from ROIs in locations where pixels darkened (Fig. 5B, green ROIs). The M1 time course confirmed that reflectance did not change from baseline until movement completion (Fig. 5C; +2.6 s from Cue). Averaging across sessions therefore washed out the positive reflectance observed after movement onset in single sessions (Figs. 3I & 4C-D). Similarly, averaging attenuated the magnitude of positive reflectance observed in the withhold condition (Fig. 3J). In PMd, averaging revealed an initial negative peak during movement (Fig. 5C; +1.0 to +1.5 s from Cue) that was not observed in single sessions (Figs. 3I & 4A-B). Nevertheless, the timing of the largest negative peak was consistent between PMd and M1. Finally, averaging showed that reflectance change in the full FOV remained near baseline across conditions (Fig. 5C). Despite differences between average time courses and the single session counterpart, both indicated that cortical activity was focal and limited to movement conditions.

### Activity maps overlap limited portions of the forelimb representations

Next, we examined the spatial relationship between domains and the motor map. For each movement condition we generated an *activity map* that flagged pixels that darkened in multiple sessions. To that end, we first applied the same t-test analysis from Fig. 3G to generate a thresholded movement frame for every condition in 8 sessions. Those frames were then co-registered across sessions and synthesized into a single frame. Every pixel was assigned a value from 0-8 to reflect the number of sessions in which it was flagged in a movement frame (e.g., Fig. 5D). To obtain an activity map, the single frame was thresholded to include only pixels that were flagged in ≥50% of sessions (Fig. 5E-F). The general organization of domains (i.e., red patches) was more similar across conditions within an animal, than for the same condition across animals. In all conditions, domains were primarily in M1 and PMd forelimb representations and were more concentrated in arm zones than in hand zones. Activity maps were co-registered with the motor maps for quantification. The most lateral domains (bottom of panels) overlapped the face representation and were excluded from analyses. Nevertheless, domains overlapped only 18% (median, IQR = 11–20%) of the M1 forelimb representation and 16% (median, IQR = 9– 23%) of the PMd forelimb representation (Fig. 5G-H). Even the activity map with the greatest surface area – precision grip in monkey G –intersected with only 18% of the M1 forelimb representation and with 38% of the PMd forelimb representation.

The relatively small activity maps prompted us to redo the analysis with a less restrictive approach. We therefore generated an activity map at every time point and quantified the cumulative overlap of those maps with the motor map (Fig. 5I-J). Indeed, cumulative overlap with the M1 forelimb representation (median = 34%, IQR = 29–43%) and the PMd forelimb representation (median = 35%, IQR = 25–59%), was twice as large as the overlap from the 5-frame average. More importantly though, the less restrictive approach confirmed that task-related activity was limited to small subzones (∼35%) of the forelimb representations. Animating the cumulative activity maps (Videos 2-3) also showed that their spatial configuration was consistent with the 5-frame activity maps.

### Clustering time courses recapitulates spatial patterns of activity

Next, we examined the spatial patterns returned form clustering time courses. The clustering algorithm considered entire time series and required no supervision. Our motivation for generating clustered maps was to evaluate whether activity patterns reported thus far from average frames, are extractable without our input on frame selection. To that end time courses were generated from a grid of cells (cell = 15×15 pixels; 0.27×0.27 mm) that covered the FOV (Fig. 6B & 6E). *k-medoids* clustering was conducted on average time courses (n=8 sessions). For comparison, an average movement frame was generated from the same time series (Fig. 6A & 6D, +4.1 to +4.6 from Cue).

**Figure 6.**
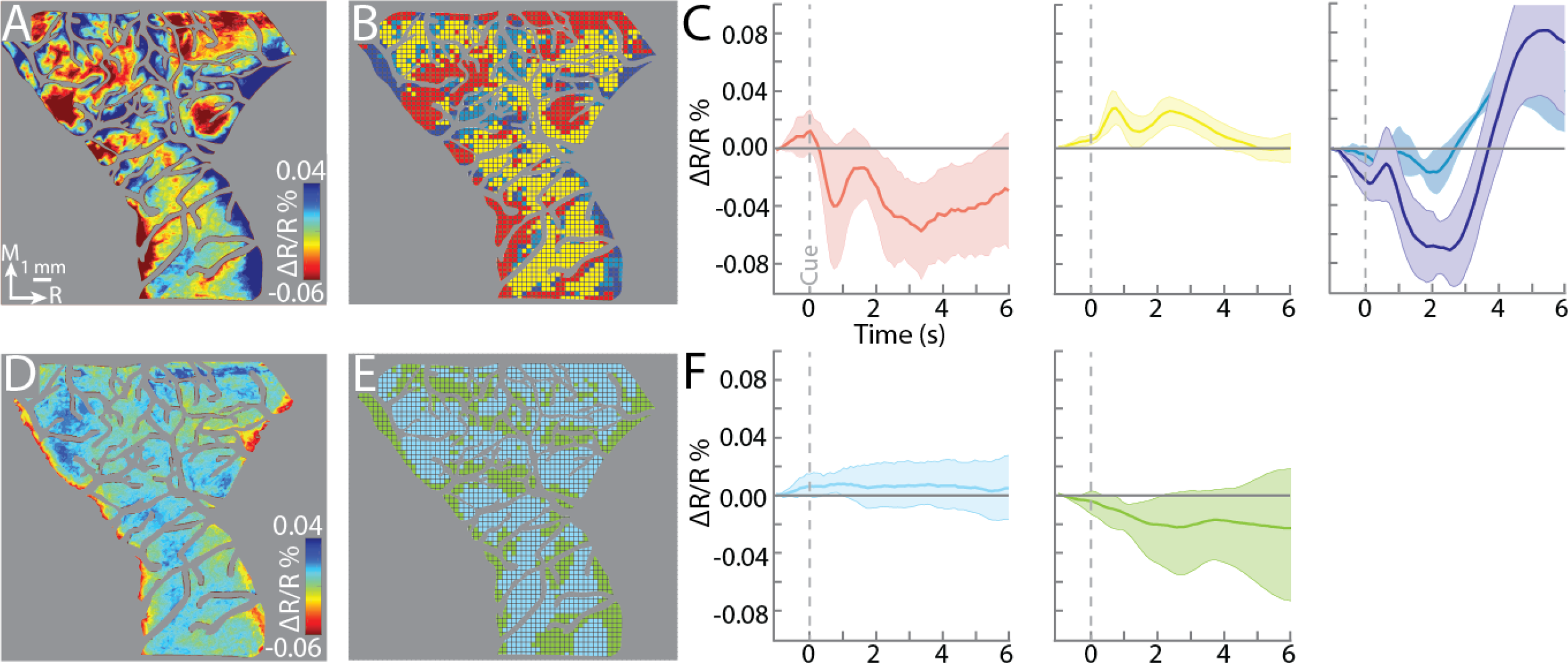
Clustering time courses recapitulates spatial patterns of activity. (A) Mean movement frame (+4.1 to +4.6 s from Cue) for the power grip condition (8 sessions, 467 trials, monkey S). Reflectance change was clipped to median ± SD pixel value across frames. Scale and intensity bars also apply to (D). (B) K-medoids identified four clusters. Each grid cell (15×15 pixels or 0.27×0.27 mm) was an ROI for time course measurement. (C) Time courses of reflectance change (mean ± SD) for each cluster. Line plot colors match clusters in (B). (D-F) Same as (A-C), but for the withhold condition (8 sessions, 488 trials).

In the precision grip condition (Fig. 6A-C), 4 clusters were found (Fig. 6B). The spatial organization of the red cluster approximated activity domains in the movement frame (Fig. 6A, hottest colors). The yellow cluster corresponded mostly with pixels that darkened at lower intensities in the movement frame (Fig. 6A, yellow and green). Most of those pixels did not meet the statistical threshold set for activity maps in Fig. 5H. The light and dark blue clusters corresponded with major blood vessels in the central sulcus, arcuate sulcus, and the precentral dimple. The time course profile of each cluster (Fig. 6C) was consistent with its spatial affiliation with the movement frame. Thus, the red cluster had a main negative peak approximately +3.5 s from Cue, which was consistent with the average time course in Fig. 5C. The yellow cluster had some positive reflectance, but it was generally near baseline as expected from pixels with no task-related reflectance change. The blue clusters had distinctively large, and delayed, positive peaks, as expected from blood vessels near active zones.

In the withhold condition (Fig. 6D-F), only 2 clusters were found (Fig. 6E). The cyan cluster (Fig. 6E) corresponded with locations of pixel brightening (Fig. 6D, blues). The green cluster corresponded with major blood vessels, which darkened at least near the central and arcuate sulci (Fig. 6D, hot colors). Average time courses confirmed that the clusters had distinguishable profiles even though their relative modulation from baseline was small (Fig. 6F). Both time courses were consistent with ones reported in Fig. 3F-J & 5C.

Observations in Fig. 6 were generally consistent in monkey G (Fig. S2). Spatial correspondence between activity maps and clustered maps was not as apparent as in Fig. 6, but it was present nonetheless for the precision grip and withhold conditions. Also, similar time course profiles were identified for the clusters in monkey G. Consistency between clustered maps and activity patterns in both monkeys, indicates that average frames reliably summarize activity patterns within a time series.

### Activity maps partially overlap across conditions

Next, we examined spatial correspondence between activity maps across conditions. Fig. 7A shows pairwise comparisons of activity maps from Fig. 5E. Teal colored pixels were present in both conditions. Other colors depict pixels that were present in 1 condition only. Comparisons to precision grip mostly returned teal and red, which means that activity maps from power grip and reach-only were a subset of the activity map from precision grip. In contrast, comparison between power grip and reach-only returned all three colors, which indicates less spatial overlap between those two conditions. Pixels outside arm or hand zones were made transparent and excluded from further analyses. Most of those pixels were in the face representation (bottom of each panel) and were likely present due to condition differences in the timing, or vigor, of anticipatory licking for reward.

**Figure 7.**
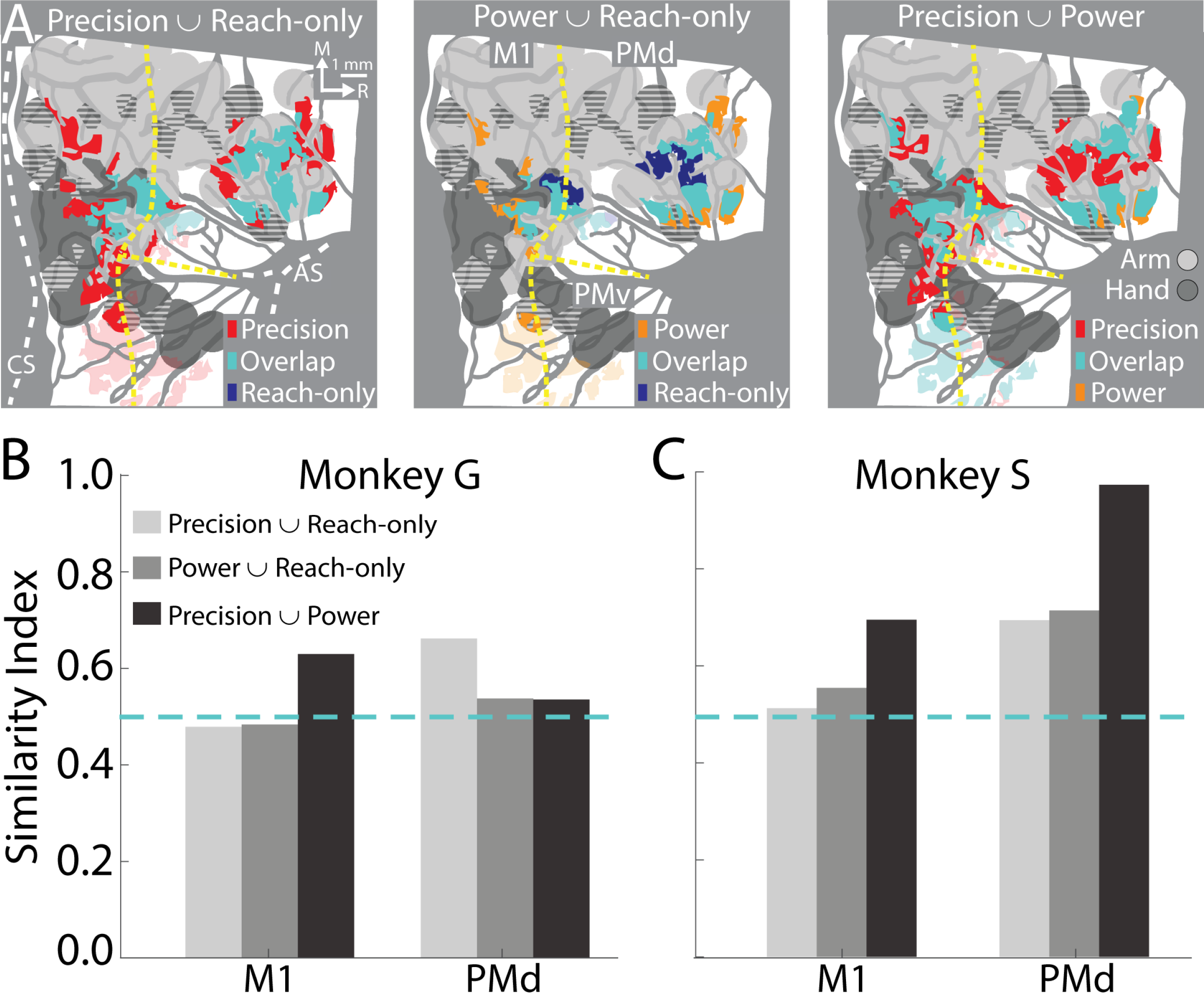
More overlap than separation between activity maps from different movement conditions. (A) Pairwise comparisons of activity maps from Fig. 5E (monkey G). Teal pixels show spatial overlap between pairs of activity maps. Red, orange, and blue pixels indicate that activity was in only one of the two activity maps. Pixels were excluded if they did not belong to a cluster that was ≥ 200 pixels^2^ (≥ 0.06 mm^2^). Transparent pixels were outside the forelimb representations. (B) Similarity index for pairwise comparisons in (A). Dotted line marks equal number of overlap pixels and condition-specific pixels. (C) Same as (B) for monkey S.

For each pair of activity maps, a similarity index (SI) was calculated from the ratio of overlap pixels and condition-specific pixels (12 comparisons: 3 conditions x 2 cortical areas x 2 monkeys). For a pair of activity maps, SI approaching 0 indicates no spatial overlap, whereas SI=1 indicates complete spatial overlap. SI values were lower in monkey G than S (Figs. 7B-C). This was mostly related to domain size differences in monkey G as opposed to spatial offsets in domains between conditions. Nevertheless, the average SI (median = 0.60, IQR = 0.54–0.71) across both monkeys showed that pairs of activity maps had more overlap than non-overlap. We conclude from this analysis that across movement conditions, domains were concentrated in the same M1 and PMd zones.

### Forelimb use differences across conditions

We examined whether forelimb use could shed light on the spatial relationship between activity maps in monkey G. Because the precision grip had the largest activity map (Fig 5G & 7A), we reasoned that forelimb use may be greater in that condition as compared to the other two movement conditions. To test that possibility, we recorded EMG activity from 7 forelimb muscles and measured kinematics for 12 degrees of freedom in the forelimb. Fig. 8A shows average profiles for 2 arm muscles and 2 hand muscles. Similarities and differences were evident across conditions even in this small sample of muscles. For example, the largest EMG peaks occurred when the hand interacted with the target object. That time point was immediately before object lift in the reach-to-grasp conditions and coincided with target contact and hold in the reach-only condition. Also, in the reach-to-grasp conditions, similar profiles were observed for triceps, ECRB, and FDS. In contrast, the deltoid peak in the precision grip distinguished it from the other two conditions. Similarly, in the reach-only condition, the FDS peak and limited ECRB activity stood out from the other conditions. Similarities and differences were also apparent in the flexion/extension profiles of arm and hand joints (Fig. 8B). For example, in the reach-to-grasp conditions, profiles were similar for the shoulder and elbow. But digit and wrist activity were protracted in the precision grip as compared to the power grip. Notably, there was almost no digit or wrist activity in the reach-only condition.

**Figure 8.**
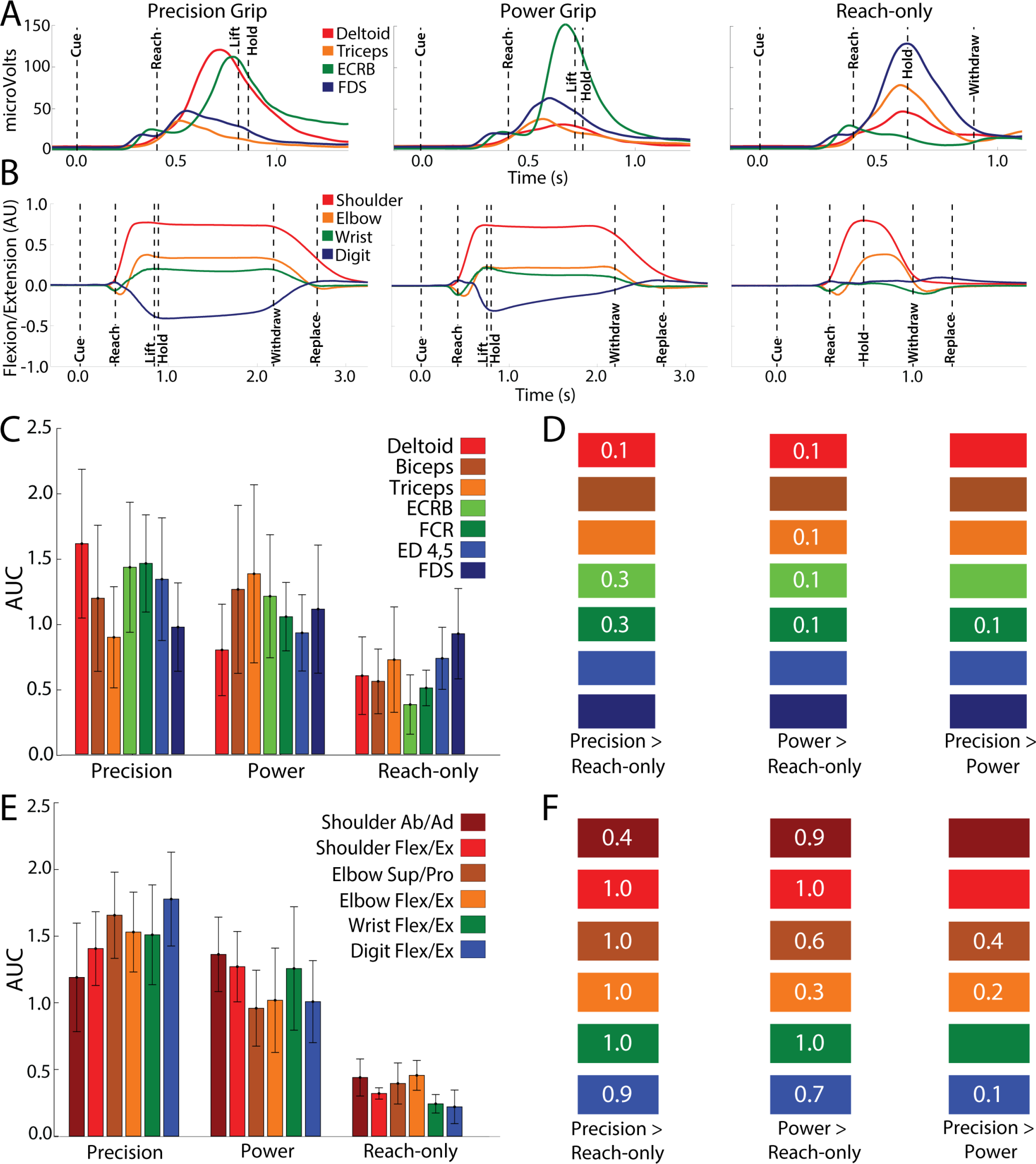
Muscle activity and joint kinematics across conditions. (A) Mean EMG activity for 2 arm muscles and 2 hand muscles (6-14 sessions, 308-595 trials, monkey G). ECRB: extensor carpi radialis brevis. FCR: flexor carpi radialis. Dashed lines mark median times of task phase onset. (B) Mean flexion/extension changes in 2 arm joints and 2 hand joints (5–15 sessions, 143– 405 trials, monkey G). Note, longer time scale here as compared to (A). (C) Area under curve (mean ± SD) for muscle activity from trials in (A). (D) Between condition comparisons (t-test) of muscle activity. Numbers are effect size estimates (Cohen’s d); empty rectangles had no effect. Colors correspond to (C). (E-F) are same as (C-D), but for joint angle kinematics.

We quantified EMG and joint profiles from the area under the curve (AUC) during Cue to Replace. AUC values were generally larger in the reach-to-grasp conditions as compared to the reach-only condition (Fig. 8C & 8E). For each muscle and joint, AUC was compared pairwise between conditions (t-test) and the effect size was quantified with Cohen’s d (Fig. 8D & 8F). Effect size is considered small at Cohen’s d of 0.2, medium at 0.5 and large at 0.8 (Cohen, 1988). Pairwise comparisons showed that wrist and arm muscles were more active in reach-to-grasp conditions as compared to the reach-only condition (Fig. 8D). The small difference between the precision and power conditions was limited to the wrist flexor. Condition differences were more apparent in joint angle kinematics than in EMG activity. Specifically, differences between reach-to-grasp and reach-only conditions were present in all joints (Fig. 8F). Smaller differences were found between the precision and power conditions, and they were mostly limited to digits and elbow (Fig. 8F). Collectively, EMG and kinematics showed that reach-to-grasp conditions involved greater forelimb activity than the reach-only condition, and that the precision grip involved greater forelimb activity than the power grip. Thus, forelimb use differences across conditions support the activity maps differences across conditions.

## Discussion

We examined the relationship between movement-related cortical activity and motor maps in motor (M1) and dorsal premotor (PMd) cortex. Intrinsic signal optical imaging (ISOI) showed that activity clustered in *domains*, which collectively overlapped small fractions of the forelimb representations. Thus, we infer that neural activity for arm and hand functions are not uniformly distributed throughout the forelimb representations. Instead, we propose a functional organization wherein subzones of the forelimb representation are preferentially tuned for actions.

### Measuring activity in M1 & PMd

Two provisions facilitated our findings. First, we used ISOI to measure cortical activity. ISOI is well-established for studying sensory cortex (Bonhoeffer and Grinvald, 1991; Chen et al., 2001; Grinvald et al., 1986; Ts’o et al., 1990), but few studies have leveraged it for cortical control of movement (Friedman et al., 2020a; Siegel et al., 2007). Functional MRI has been used more extensively to investigate reaching and grasping in monkeys and humans (Cavina-Pratesi et al., 2010; Gallivan and Culham, 2015; Nelissen and Vanduffel, 2011). Our objective, however, was better served with the higher contrast and spatial resolution afforded from ISOI. Similarly, calcium imaging has gained traction in cortical control of movement because of temporal fidelity to neural spiking. But in monkeys, the field-of-view (FOV) in mini-scopes and 2-photon microscopes, captures only a small zone of one forelimb representation (Bollimunta et al., 2021; Ebina et al., 2018; Kondo et al., 2018). In contrast, the present FOV was large enough for both M1 and PMd forelimb representations. Second, we obtained motor maps from the same chronic chambers and used them to quantify the spatial organization of the activity maps. Motor maps also confirmed that the forelimb representations were in the FOV in every imaging session. Thus, conducting both ISOI and motor mapping was central to our results and interpretation.

### Movement activates punctate domains

Activity maps overlapped small fractions (∼16%) of the M1 and PMd forelimb representations. Our criteria for activity maps flagged pixels that darkened significantly in ≥50% of imaging sessions. Thus, the constituent domains of activity maps were not exclusive zones of task-related activity. Instead, domains reported locations where task-related activity was both robust and repeatable across sessions.

We consider explanations for the relatively small size of the activity maps. It is unlikely that ISOI lacked sensitivity for capturing the full extent of cortical activity and thereby underreported activity map sizes. If anything, ISOI sensitivity to subthreshold electrophysiological signals implies overestimation of the spatial extent of modulated neural activity (Frostig et al., 2017; Grinvald et al., 1994), which is remedied by focusing on early segments of the ISOI response. With this strategy, ISOI reports cortical organization and connectivity at columnar resolution (Bonhoeffer and Grinvald, 1991; Card and Gharbawie, 2022, 2020; Friedman et al., 2020b; Lu and Roe, 2007; Vanzetta et al., 2004). Extensive task training, as done here (∼2 years), refines network activity and reduces metabolic demand in M1 (Peters et al., 2014; Picard et al., 2013). Both factors could have shrunk activity maps from larger sizes in earlier training stages. Importantly, the relatively small size of activity maps reveals an organizational aspect of M1 and PMd. Specifically, peak neural activity for reaching and grasping is concentrated in focal zones within the forelimb representation. It stands to reason that domain-free zones within the forelimb representation, could be preferentially tuned for other arm and hand actions. This notion of functional clustering in M1 and PMd has support from long-train ICMS (500 ms), which maps multi-joint actions (e.g., reach, manipulate, climb) to contiguous zones in M1 and PMd (Baldwin et al., 2016; Gharbawie et al., 2011; Graziano et al., 2002; Kaas et al., 2013). Imaging movement-related cortical activity circumvents potential complications from interpreting movement evoked with long-train ICMS (Griffin et al., 2014). Thus, it would be instructive to investigate the functional organization of M1 and PMd with ISOI in monkeys performing a range of arm and hand tasks.

### Differentiating between domains

Whether domains of an activity map were functionally differentiable requires some interpretation. The temporal resolution of intrinsic signals is not sufficient for relating reflectance change to behavioral events that occur in close succession (e.g., reach and grasp). We are currently conducting high-density electrophysiological recordings to interrogate relationships between spiking activity in the present domains and movement. Our report on a small sample of single units recorded in other hemispheres, suggested functional differentiation between domains (Friedman et al., 2020a). Specifically, we found that a medial M1 domain was enriched with neurons that were preferentially tuned for reaching, whereas a lateral M1 domain was enriched with neurons that were more strongly tuned for grasping. Other recordings, however, from caudal M1 (central sulcus) or rostral M1 (precentral gyrus) reported spatially co-extensive encoding of reaching and grasping (Rouse and Schieber, 2016; Saleh et al., 2012; Vargas-Irwin et al., 2010). Thus, the collective evidence from the temporal dimension seems equivocal on functional differences between domains.

The spatial organization of the domains, however, is arguably more suggestive of functional differentiation. Domains that overlapped arm zones could have been tuned to different task phases than domains that overlapped hand zones. Other studies have inferred domain functions from contrasting activity maps generated from different actions. For example, Nelissen & Vanduffel (2011) identified grasp domains by comparing fMRI activity in reach-to-grasp conditions to a reach-only condition. Similar comparisons in our data showed that activity maps were more similar across conditions. This discrepancy may mean that in our task, forelimb use differences across conditions, were not large enough. We therefore cannot infer domain functions from comparing condition maps, which is consistent with other studies that used univariate analyses on fMRI measurements (Fiave et al., 2018; Gallivan et al., 2011; Nelissen et al., 2018). Nevertheless, multi-variate analyses (e.g., multi-voxel pattern analysis) on the same data showed that condition identities were embedded in the spatial dimensions of activity maps.

### Distinct time courses in M1 and PMd

ISOI time courses were sensitive to functional differences between M1 and PMd. Specifically, the early dark peak in the average PMd time course was absent from M1. That peak is consistent with preparatory activity characteristic of PMd (Crammond and Kalaska, 1996; Weinrich and Wise, 1982). Our task enforced a 5-second delay between trial initiation and Go Cue. That delay likely separated ISOI signals for movement preparation from those for movement execution, thereby revealing successive peaks in the PMd time course. In the *observation* experiment, activity was absent in M1 but present in PMd, which is consistent with previous findings and higher concentrations of mirror neurons in premotor areas than in M1 (Grèzes et al., 2003; Papadourakis and Raos, 2019; Raos et al., 2007). Finally, in the Withhold condition, M1 activity remained near baseline, whereas positive reflectance was evident in the PMd time course. The results collectively show that PMd activity could have been related to motor cognitive process as well as movement, whereas M1 activity was more closely related to movement.

### Main negative peak locked to movement

Pixel darkening peaked several seconds after movement onset. Our time courses from single sessions were consistent with those reported in the few studies that used ISOI with forelimb tasks (Friedman et al., 2020a; Heider et al., 2010). Our three control experiments confirmed that the relatively slow negative peaks were predicated on movement execution. Nevertheless, the lag between movement onset and peak pixel darkening was 2-3 times longer than expected from stimulus onset (peripheral or ICMS) and peak darkening in sensorimotor cortex (Card and Gharbawie, 2020; Chen et al., 2001; Friedman et al., 2020b). Slower intrinsic signals have been reported in other behavioral paradigms, which raises the possibility of time course differences between awake and anesthetized ISOI (Grinvald et al., 1991; Tanigawa et al., 2010). Fundamental differences could also exist between intrinsic signals evoked from movement (active) and those from stimulation (passive). In our task, the arm and hand were active for 1-2 s per trial. In contrast, sensory stimuli are typically confined in both space and time to optimize focal activation in cortex (e.g., S1 barrel, V1 orientation column).

The protracted peak darkening should be considered in relation to the triphasic time course of reflectance change in ISOI (Chen-Bee et al., 2007; Sirotin et al., 2009). Studies with stimulus evoked responses typically focus on the *initial dip*, which is the first dark peak and occurs 1-2 s after stimulus onset. The subsequent *rebound* (bright peak) and *undershoot* (second dark peak) unfold sequentially over several seconds. In our time courses, the timing of the darkest peak would have coincided with the *rebound* in stimulus-evoked responses. We offer three explanations for the timing of our dark peak. (1) Our dark peaks could have been *initial dips* locked to movement onset. In this interpretation, our movement-evoked time courses would be slower iterations of stimulus-evoked responses. (2) Our dark peaks could have been *initial dips* locked to movement offset. Directly testing this possibility requires systematic shifting of movement offset times, which we did not explore. (3) Our dark peaks could have been the *undershoot* locked to movement onset. In this interpretation, our movement-evoked time courses would be consistent with stimulus-evoked time courses. The average PMd time course may support this interpretation as it had a small negative peak followed by a larger negative peak. In M1, the *initial dip* may have been undetected because it was too small, obscured by motion artifact, or both.

## Conclusion

We showed that in M1 and PMd, activity related to reaching and grasping was concentrated in subzones of the arm and hand zones. Thus, the spatial dimension of the forelimb representation is central to its functional organization. Taking this feature into consideration in electrophysiological studies is likely critical to understanding relationships between the recorded signals and movement control.

## Acknowledgments

Project was supported with funds from NIH (R01 NS105697); Whitehall Foundation (2017-12-94), and University of Pittsburgh Brain Institute. We are grateful to ToniAnn Zullo for outstanding animal care before, after, and during, all procedures. We thank Nick Card for advice on analyses.

## Contributions

NGC & OAG designed and performed experiments, analyzed data, and wrote the manuscript. NGC prepared all figures. OAG obtained funding.

## Conflict of interest

none.

---

**Supplementary Figure 1.**
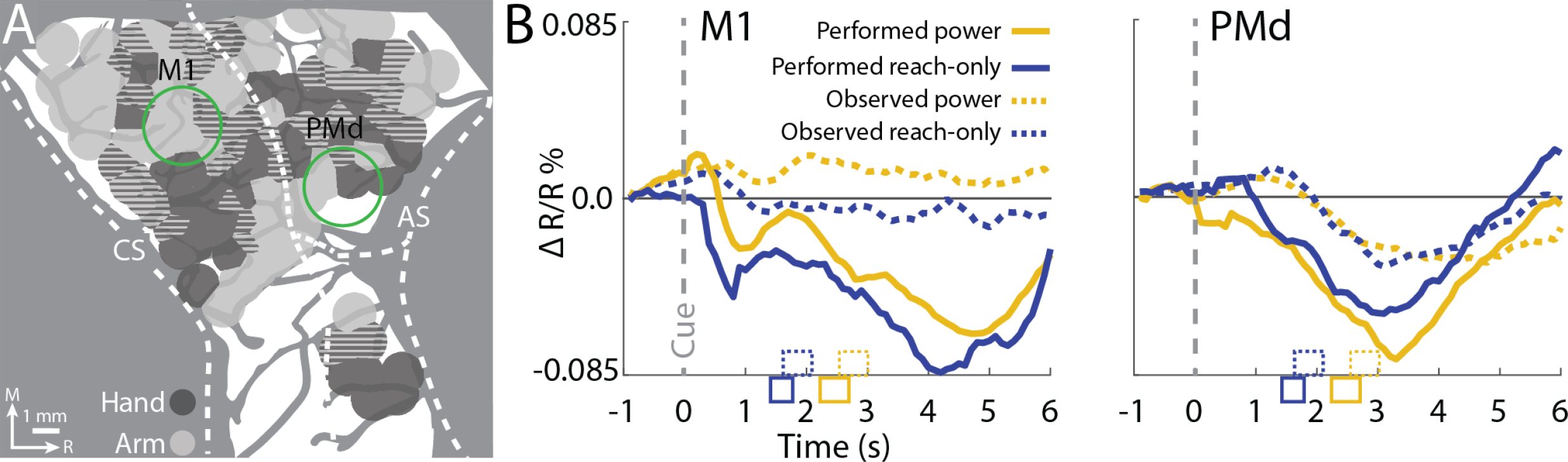
Movement observation drives reflectance change in PMd but not M1. (A) Green ROIs in M1 (2.7 mm diameter), PMd, and full FOV (monkey S). (B) Time courses from 2 movement execution conditions (35 trials/condition) and 2 movement observation conditions (81 trials/condition). Makers on x-axis indicate time (mean ± SD) of movement offset. Markers match line plot formats.

**Supplementary Figure 2.**
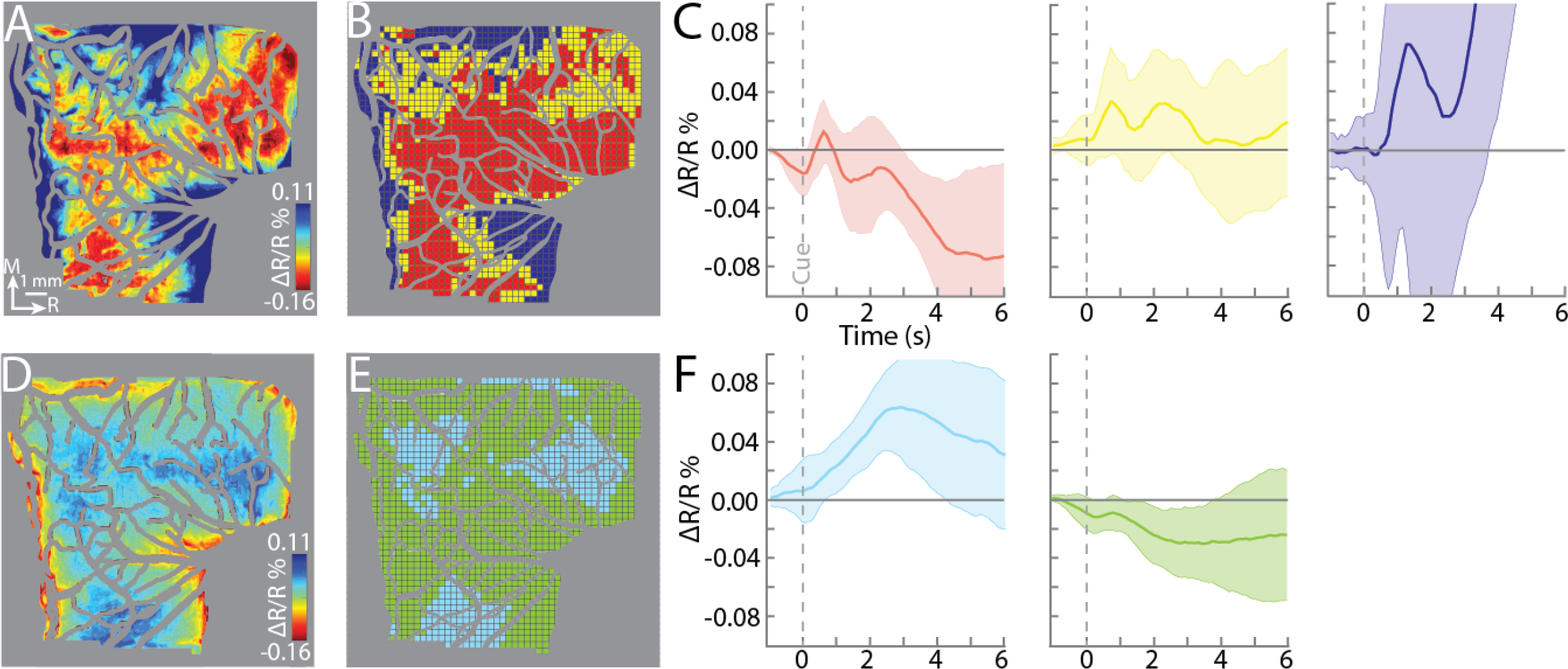
Time course clustering recapitulates spatial patterns of activity. (A) Mean movement frame (+4.1 to +4.6 seconds from Cue) for the precision grip condition (8 sessions, 404 trials, monkey G). Reflectance change was clipped to median ± SD pixel value across frames. Scale and intensity bars also apply to (D). (B) Three clusters were identified with k-medoids. Each grid cell (15×15 pixels or 0.27 x 0.27 mm) was an ROI for time course measurement. (C) Time course of reflectance change (mean ± SD) for each cluster. Line plot colors match clusters in (B). (D-F) Same as (A-C), but for the withhold condition (8 conditions, 376 trials, monkey G).

**Video 1. Pixel darkening in ISOI lags movement.** *Left:* representative trial from the precision grip condition filmed from overhead (monkey G). Task was performed with left forelimb and right forelimb was restrained. Video plays at half-speed. Time in bottom right is relative to start of ISOI acquisition; Cue onset was at 1.0 s. Movement started at 1.3 s and was complete by 3.6 s. *Right:* Average ISOI time series (8 sessions, 404 trials) from the same condition. Beyond the image processing described in the methods, frames were temporally smoothed with a 5-frame sliding window. Brown outline flags clusters of pixels that darkened beyond −0.08 (∼40^th^ percentile). Note, that domains did not form until movement completion.

**Video 2. Spatio-temporal development of cumulative activity maps.** A 5-frame moving average was calculated at each time point and compared to the baseline frame (monkey G). Pixels flagged for darkening in 1 frame (t-test, p<0.0001) remained red for the rest of the animation (i.e., cumulative activity map). The activity map for the reach-to-grasp condition was considerably larger than for the reach-only condition. Nevertheless, neither map filled the forelimb representation. The ON pulse denotes the movement phase of an average trial. Animation plays at half speed.

**Video 3. Spatio-temporal development of cumulative activity maps.** Same as video 2, but for monkey S. Note similar map size between conditions.

## Notes

### Competing Interest Statement

The authors have declared no competing interest.

